# Higher attention is associated with stronger sensorimotor connectivity during a target pursuit task

**DOI:** 10.1101/2025.07.31.667972

**Authors:** Andrew Paek, Shikha Prashad

## Abstract

Attention is critical in processes related to motor learning and recovery. While neuroimaging studies have highlighted its relevance to frontal-parietal networks, it is unclear how attention affects movement-related activity from the sensorimotor regions of the brain during a motor task. In this study, participants pursued a moving target with a computer mouse, and the level of attention was manipulated via two conditions in which the target moved in either a predictable or unpredictable fashion. We compared event-related desynchronization (ERD) between rest and movement as well as coherence during movement between the predictable and unpredictable trials. We found that alpha- and beta-band ERDs in the contralateral central areas did not change significantly between the two conditions. However, unpredictable trials had larger alpha-band suppression in the frontal and parietal areas, larger beta-band suppression in the ipsilateral parietal area, and larger alpha-band functional connectivity across the central areas. Our study highlights that performing limb movements with higher levels of attention was associated with stronger cortical communication in the sensorimotor areas, rather than strengthening neural activity in these areas. However, higher levels of attention were associated with stronger activation in the frontal and parietal areas which reflects engagement in attention-related networks. This increased activation of attentional networks and stronger sensorimotor communication may facilitate the formation of neural connections that occur in motor skill training and recovery.

**Impact Statement:** How the brain’s attentional networks interact with motor learning processes is still poorly understood. In our study, we found that performing motor tasks with higher levels of attention was associated with stronger frontal and parietal ERDs, and stronger bilateral communication in the central motor areas. This corroborates with prevailing theories that the prefrontal cortex, which moderates attention, induces communication within the sensorimotor areas to promote the formation of neural connections that occur with motor learning.

## Introduction

Attention is critical for the learning of motor skills. This has been demonstrated in behavioral studies, where dividing one’s attention with a secondary cognitive task has been shown to compromise motor learning (Kahneman, 1973; Song, 2019; Wickens, 2020). In addition, attention aids motor recovery in stroke rehabilitation. For example, recent studies highlight how motor recovery is substantially higher when a stroke patient performs exercises with active engagement, as opposed to exercises where the patient passively experiences limb movements that are carried out with significant assistance from the therapist (Hogan et al., 2006; Krebs et al., 2009; Venkatakrishnan et al., 2014). These findings have led to efforts to design neuroimaging systems that can monitor a stroke survivor’s level of engagement during rehabilitative exercises (Bartur et al., 2017, 2020).

These behavioral studies demonstrate that attention can facilitate motor learning, but the neural mechanisms behind this process are unclear. Current neuroimaging studies have highlighted the control of attention primarily in neural networks between the frontal and parietal areas (Corbetta & Shulman, 2002; Miller & Buschman, 2013). Scalp EEG activity from these areas has been used to formulate metrics that measure levels of task engagement and cognitive workload (Berka et al., 2007; Pope et al., 1995). These EEG-based measures become stronger when individuals perform more difficult motor tasks (Jaquess et al., 2018) and when practice schedules are varied between sessions (Lelis-Torres et al., 2017). While these studies demonstrate how frontal-parietal networks are modulated with task difficulty, there is a gap in understanding how this network interacts with sensorimotor areas during skill acquisition.

We propose that greater levels of attention may lead to more recruitment of motor-related cortical areas. Steady-state somatosensory-evoked potentials are known to be modulated by the participant’s selective attention (Ahn et al., 2016; Giabbiconi et al., 2004; Yao et al., 2013). These rhythms are observed when different parts of the body are stimulated using tactors that vibrate at different frequencies. When a participant selectively focuses their attention on a particular body part receiving such stimulation, brain waves in the sensorimotor areas become attuned to the corresponding frequency of the vibration (Ahn et al., 2016; Giabbiconi et al., 2004; Yao et al., 2013). Just as attention can modulate sensory-related cortical oscillations, we suspect that attentional processes can also modulate movement-related cortical activity. This modulation may be associated with higher levels of cortical engagement that can facilitate motor learning. Other studies have found that attention modulated cortical activity in sensorimotor areas when experimenters induced varied levels of attention by changing the difficulty of the task (Dujardin et al., 1993; Mizelle et al., 2010a, 2010b). However, the observed changes in neural activity may reflect cortical processes that respond specifically to the experimental manipulations, rather than attentional modulation. For example, the task used by Dujardin et al. (1993) involved distinguishing which one of two words presented to the participant belonged to a list of memorized words. Higher levels of attention were induced by presenting words that were closer in meaning, which may have invoked cortical activity related to semantic meaning and memory recall. Mizelle et al. (2010a, 2010b) instructed participants to aim at a target with their leg and induced higher levels of attention by making the target smaller or by connecting a weight to their leg. Such experimental conditions could have engaged cortical processes associated with visual and proprioceptive processing. In both studies, it is unclear if the changes in the observed brain activity can be attributed to attention or recruitment of other cortical areas necessary to perform the task under specific experimental conditions.

In this study, we assessed if movement-related cortical activity changed solely due to changes in attentional demand. We implemented a behavioral task where participants used a computer mouse to pursue a moving target on the screen. Attentional demand was manipulated through two conditions: 1) where the target moved in a predictable pattern and 2) where the target changed directions unpredictably. We hypothesized that pursuing a target with unpredictable movements would require more attention from participants (without changing the difficulty of the task) and result in stronger neural modulations associated with movement. We focused on movement-related event related desynchronizations (ERDs) that were recorded with scalp EEG. ERDs are characterized by a reduction in alpha- and beta-band frequencies in central areas of the scalp when individuals shift from rest to movement. ERDs are indicative of cortical engagement (Pfurtscheller, 2006; Pfurtscheller et al., 1996; Pfurtscheller & Lopes da Silva, 1999). This feature has been studied heavily with hand movements and is known to modulate in strength based on factors such as movement speed (Yuan et al., 2010), complexity (Bian et al., 2018), and grip commands (Cassim et al., 2000; Nakayashiki et al., 2014). We hypothesized that when the task demanded more attention, the ERDs generated in the sensorimotor areas would become stronger. We also calculated coherence as a measure of functional connectivity, as we anticipated that higher attentional demand would increase communication from sensorimotor areas to other regions in the brain.

## Methods

### Experimental Hardware

Scalp EEG was recorded with 64 channels at a rate of 1000 Hz with an active electrode system (Brain Products, ActiChamp, Gilching Germany). Two channels were used to record horizontal and vertical electrooculography activity by placing sensors on the left temple and just under the right eye. The ground and reference electrodes were placed respectively on the left and right ear lobes. Visual stimuli were displayed on a 24-inch monitor (Dell S2417DG) at a rate of 144 Hz. Visual cues for the task were presented in a customized protocol that was programmed with Presentation (Neurobehavioral Systems Inc., Berkeley, CA). This software was also used to log the timing of the events for each trial, the position of a moving target that was displayed to participants, the position of a cursor that participants controlled with a computer mouse, and when participants pressed the space bar key on a keyboard.

### Behavioral Task

Twenty able-bodied participants (15 female, age: 20.95 ± 4.59 years) volunteered for the study. All participants were right-handed (laterality quotient mean: 94.26 ± 11.93, minimum: 66.67). They provided written consent as approved by the Washington State University Institutional Review Board. After they provided consent, they filled out a written survey that inquired their age, gender, race, and handedness through the short form of the Edinburgh Handedness Inventory (Veale, 2013).

At the beginning of the experiment, the participant donned a cap with EEG electrodes. Electrolyte gel was applied to reduce the impedance between the electrode and the scalp to less than 50kOhms. We collected baseline data where participants focused on a white crosshair for one minute followed by resting with their eyes closed for one minute. This baseline was collected before and after the behavioral task.

For the behavioral task, participants were seated next to a table with a desktop computer monitor that displayed visual stimuli. The participants were also provided with a computer mouse that they controlled with their right hand and a computer keyboard which they could press with their left middle and ring finger placed on the space bar. The screen displayed a white horizontal track with a yellow circle as the target and a smaller blue circle as the cursor as shown in Figure 1A. Participants controlled the blue circle with the computer mouse while the yellow target moved along the track.

**Figure 1).**
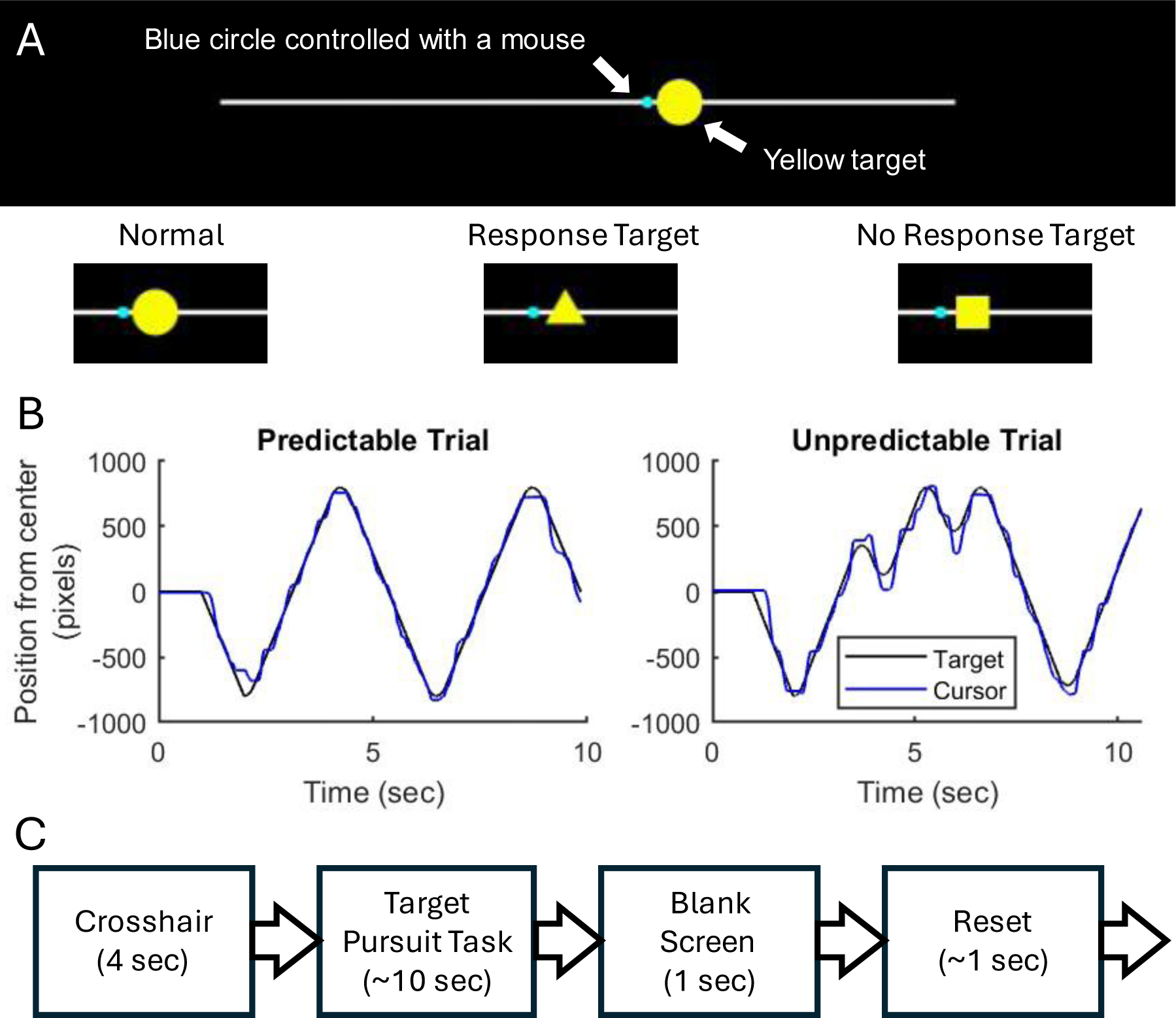
Behavioral Task Design. Participants controlled a blue cursor to follow a yellow target on a white track as shown in (A). For most of the time in each trial, the target was a circle, but could change to a triangle or square as shown. Participants were instructed to press the space bar when the target changed to a triangle and omit key presses when the target changed to a square. (B) shows examples of the position of the target (in black) and the cursor (in blue) from a single exemplary trial. (C) shows a flow chart of the events that occurred for each trial.

As the target moved, participants were instructed to keep the blue circle as close to the center of the yellow target as possible. In each trial, the target started at the center of the bar and either moved left or right. When the target reached either end of the track, it changed direction so that the target moved back towards the center. The target was programmed to travel four times the distance of the horizontal track, and at a speed where it would take approximately two seconds to travel from one end to the other. The target was also programmed such that changes in direction were dampened to avoid abrupt changes in cursor motion. Trials were approximately 10 seconds long in duration. Figure 1B shows representative examples of the target and cursor position from a trial in each condition.

The yellow target moved along the horizontal track in two task conditions: predictable and unpredictable. In the predictable condition, the target moved in a simple pattern where it only changed directions once it reached either end of the horizontal track. In the unpredictable condition, the target changed direction 3-5 times in the middle of the track. This called for the participant to maintain their attention on the target. The times when the target changed direction was designed to not occur too early or too late into the trial, or too frequently. This was programmed such that no directional changes occurred until the target travelled half of the track length or in the end when it only had half of the track length remaining to travel. When an unexpected direction change occurred, the next unexpected direction change only occurred when the target traveled a tenth of the track distance. Within these constraints, the timing and frequency (3-5 times) of the direction changes were randomized for each trial.

While participants pursued the moving target, they also performed another simultaneous task to measure their attention. This task was based on the Sustained Attention to Response Task (SART) (Robertson et al., 1997), where individuals are presented with a rapid series of numbers that appear briefly, and instructed to respond as quickly as possible by pressing a key when a number appears. They are also instructed to omit key presses only when a particular digit appeared (i.e., press the key as early possible for any number except when number “3” is shown). In this study, participants were instructed to monitor the moving target’s shape and respond to changes in the shape by pressing the space bar on the keyboard with their left hand. When the target changed from a circle to a triangle, the participants were instructed to press the space bar as quickly as possible. When the target changed from a circle to a square, the participants were instructed to omit pressing the space bar. Throughout a block of trials, the square had a 25% chance of appearing while the triangle had a 75% chance of appearing. For each trial, the target’s shape changed four times. The sequence of target shapes was randomized throughout the block of trials, such that it was possible for a trial’s shape change sequence to have all triangles or multiple squares. The behavioral task was designed such that the shape changes in the target did not occur too early or late within a trial, or too rapidly. This was programmed such that shape changes only occurred 1.5 to 9 seconds after the trial began with the target presentation, and the minimum time duration between each shape change was set to 1.15 seconds. Within these constraints, the timing of the shape changes was randomized for each trial.

Each trial consisted of a rest period, then the target pursuit task, followed by a reset period, as shown in Figure 1C. During the rest period, participants relaxed and viewed a white crosshair for 4 seconds. Participants were also instructed to avoid moving the computer mouse during this rest period. Then the participant performed the target pursuit task as described above. At the end of the target pursuit period, a blank screen with a 1 second duration was shown. This was followed by a reset period where the participant was instructed to move the cursor back to the center of the screen that contained a circular blue target. This target was also surrounded by an unfilled light blue circle, which dynamically reduced in size to indicate when the participant correctly positioned the cursor to the center of the target. When the participant was able to maintain the cursor at the center of the track for 1 second, the next trial started with the rest period containing the white crosshair.

During the target pursuit task, the cursor’s movements were restricted along the horizontal track, regardless of the computer mouse’s vertical position. The true position of the cursor was shown during the reset period, where the cursor’s horizontal and vertical position was displayed. It was common for the cursor’s true position to drift away from the horizontal track, so this reset step helped maintain the cursor’s vertical position throughout the experiment. If the participant moved away from the center of the screen during the rest period, the reset period was called again to guide the cursor back to the center position. This was followed by another rest period.

Each participant performed 125 trials of the behavioral task. The first five trials were provided as practice to ensure that the participants understood the task. Additional instruction was provided during the practice trials to clarify the task, if needed. Then participants performed four blocks of 30 trials each. The target moved predictably in the first and third block, and unpredictably in the second and fourth block.

### Kinematic Analysis

For each participant, we calculated the accuracy of omitting a key press when the target changed into a square. First, we aggregated all the instances when the target changed into a square for all trials in each condition. A correct omission was logged when a key press was not present between the time when the target changed to a square and the time when the next shape change occurred or when the trial ended. The accuracy was calculated as the number of correct omissions divided by the total number of times the target changed into a square.

We also calculated the delay between the time when the participant pressed the space bar and the time when the target changed into a triangle. These delay times were averaged from all instances in each condition for each participant.

We calculated the distance between the target’s center and the cursor’s center as a measure of error in tracking the target. For each trial, the root mean square (RMS) of the distance in pixels was calculated. These RMS error estimates were aggregated from all trials in each condition for each participant.

For each of these measures, a paired t-test was used to compare the values between the predictable and unpredictable conditions.

### EEG analysis

#### Artifact Removal

EEG signals were processed to reduce the presence of artifacts in the recordings. First, the channels were rereferenced to the linked earlobes. Next a 4^th^ order Butterworth zero-phase high pass filter with a 0.1 Hz cutoff was used to remove the slow temporal drift in the signals. Next, artifact subspace reconstruction was used to suppress brief deflections in the signals. This used the “pop_clean_rawdata” function from the EEGLAB toolbox. We reconstructed 0.5 sec segments of data that had amplitudes which were 60 standard deviations larger compared to the portions of data with minimal deflections. Next, independent component analysis (ICA) was used to identify artifacts associated with eye blinks, muscular artifacts, heart activity, and power line noise. These were identified with the “ICLabel” functions in the EEGLAB toolbox (Delorme & Makeig, 2004). Components that had over 90% confidence in the identification were removed from the EEG data.

Since we intended to analyze functional connectivity, we utilized a bipolar montage to help reduce the likelihood of spurious connectivity values (Bastos & Schoffelen, 2016). The bipolar montage is shown in Figure 2A. This montage was used in the rest of the analysis.

**Figure 2).**
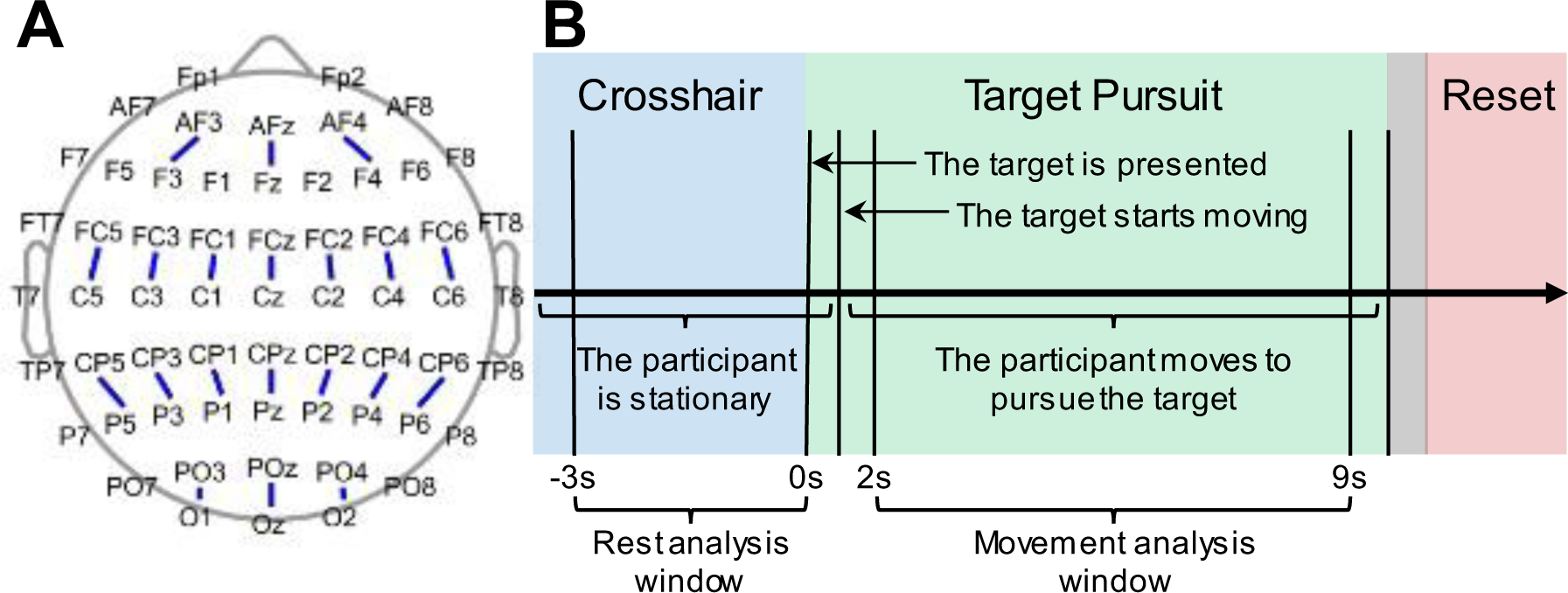
Selected sensors and time windows used for analysis. (A) The sensors used to create the bipolar montage as indicated by blue lines between pairs of sensors. (B) The timeline of a single trial, where each colored region corresponds to the periods of the trials (blue for the crosshair, green for the target, grey for the blank screen, and red for the reset). The zeroth second marker corresponds to the time when the target is presented to the participant. The braces below the timeline indicate time windows that were used to calculate the ERDs and the coherence.

#### Spectral Analysis

For each bipolar channel, we extracted the spectral power in the alpha- and beta-bands. First, 1-second time windows with an overlap of 900 milliseconds were extracted. For each time window, the EEG data were multiplied with a Hamming window to reduce spectral leakage. The power spectra were then extracted using the Fast Fourier Transform with an FFT length of 2048 samples. Based on the power spectra, the power was extracted in the alpha (8-13 Hz) and beta (20-30 Hz) bands by calculating the sum of the power spectral values in the respective frequency bins.

For each trial, the power in each of these bands was extracted from the rest period (when the participant viewed the white crosshair) and the movement period (when the participant moved the cursor to pursue the target). Within each period, power was estimated as the averaged power across the 1-second time windows that occurred during each period. The ERD was calculated in each trial as the ratio between the power during the movement period over the rest period (movement / rest). These movement and rest periods are shown in Figure 2B. The rest period was designated as the 3-second period when the crosshair was shown just before the target was presented. The movement period was designated as the 7-second period that occurred 2 seconds after the target was presented. For each participant, the average ERD was calculated across trials for each condition (predictable vs. unpredictable), and the base-10 logarithm of the average ERD was calculated for each participant. The paired sample t-test was used to compare these power values between the two conditions. To observe the spatial topography of the ERDs, scalp maps of the average ERD across participants were plotted.

#### Coherence

We explored if different levels of attention affected functional connectivity between the contralateral sensorimotor region to other areas of the brain during target pursuit. Coherence was calculated between the C3-FC3 channel and the other bipolar channels. For each trial, coherence was calculated in the movement periods, which spanned from 2-9 seconds after the target was presented, as shown in Figure 2B, using the “mscohere” function in MATLAB (Mathworks Inc., Natick, MA). This function calculated coherence based on the averaged cross spectra from segmented time windows. A 1-second time window with an overlap of 900 milliseconds was used, with an FFT length of 2048 samples. The coherence values from the trials were aggregated based on the condition to which they belonged (i.e., predictable or unpredictable). For each participant, the coherence values were averaged across trials within their respective conditions. The paired-sample t-test was then used to compare the coherence values between unpredictable and predictable trials.

## Results

### Behavioral Results

Shown in Figure 3 are boxplots of the behavioral measures. Participants were significantly more accurate in omitting key presses when the square target was shown during the unpredictable condition (mean: 94.42% ± 6.29%) than in the unpredictable condition (mean: 89.33% ± 7.81%, p < 0.001). When the target changed to a triangle, participants were significantly slower in pressing the space bar in the unpredictable condition (mean: 433.08 ± 56.69 ms) compared to the predictable condition (mean: 409.74 ± 55.50 ms, p < 0.001). Participants also had greater difficulty keeping the cursor close to the target in the unpredictable condition, where the distance between the cursor and the target was significantly larger in the unpredictable condition (mean: 66.10 ± 11.85 pixels) than in the predictable condition (mean: 49.13 ± 8.71 pixels, p < 0.001).

**Figure 3).**
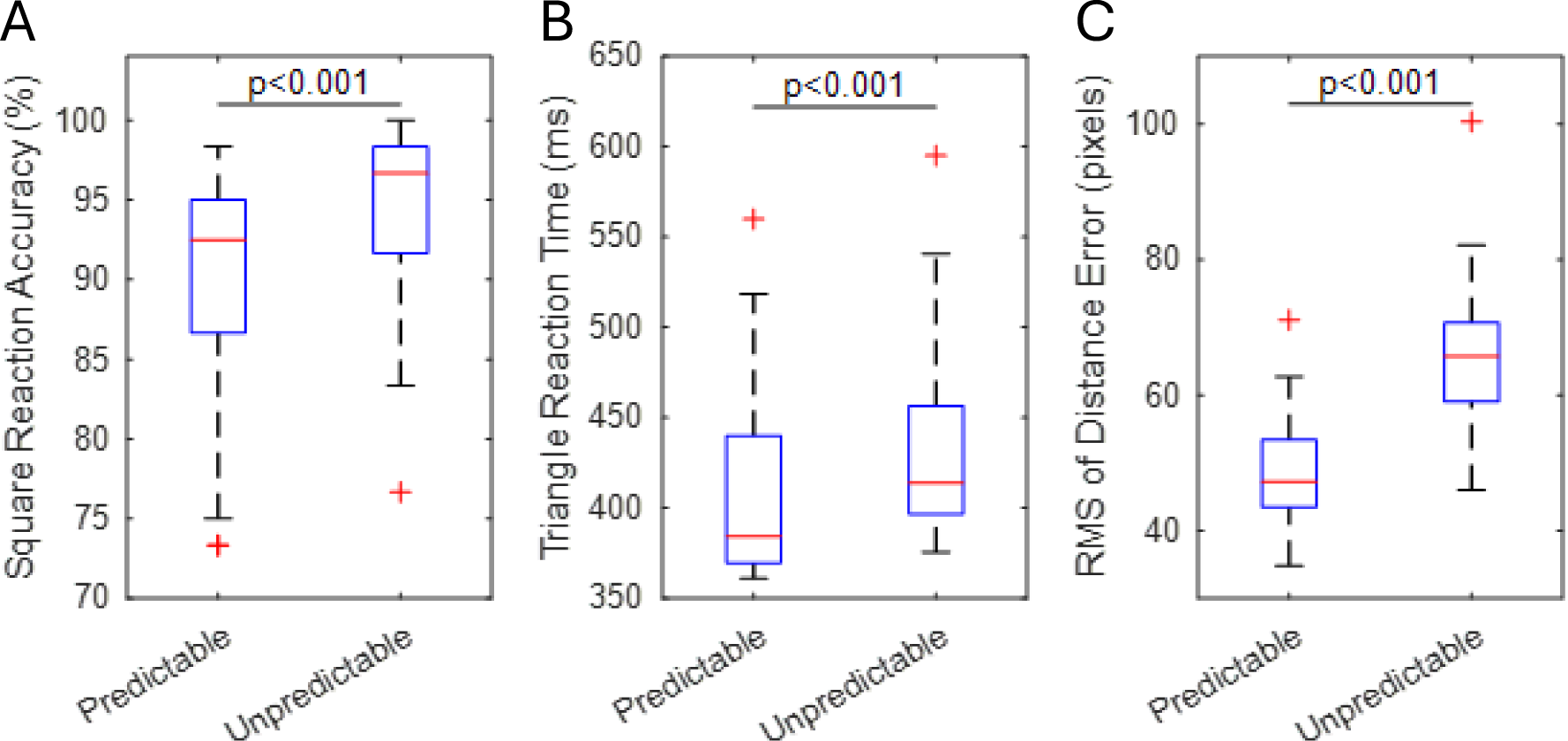
Behavioral Task Metrics. (A) shows the accuracy in omitting the key press when the target changed to a square. (B) shows the reaction time when the target changed to a triangle. (C) shows the root mean square error between the cursor and the target. Outliers are shown as red pluses and the bar indicates significance between the trials in the predictable and unpredictable conditions (p < 0.001) based on the paired sample t-test (N=20 participants).

### EEG Results

Shown in Figure 4A are the scalp maps depicting ERDs due to movement. We found that alpha-band power had notable suppression in the contralateral central areas and the posterior areas while beta-band power had notable suppression mainly in the contralateral central area. While this spatial topography was found in both conditions, the ERDs were found to be significantly stronger in the unpredictable condition. For the alpha-band, the sensors AF3-F3 (p = 0.022), AFz-Fz (p = 0.008), CP3-P3 (p=0.023), CP1-P1 (p = 0.022), and CP6-P6 (p = 0.026) had significantly stronger ERDs in the unpredictable condition compared to the predictable condition. For the beta-band, sensors CP2-P2 (p = 0.030), CP4-P4 (p = 0.003), and CP6-P6 (p = 0.039) had significantly stronger ERDs in the unpredictable condition than the predictable condition.

**Figure 4).**
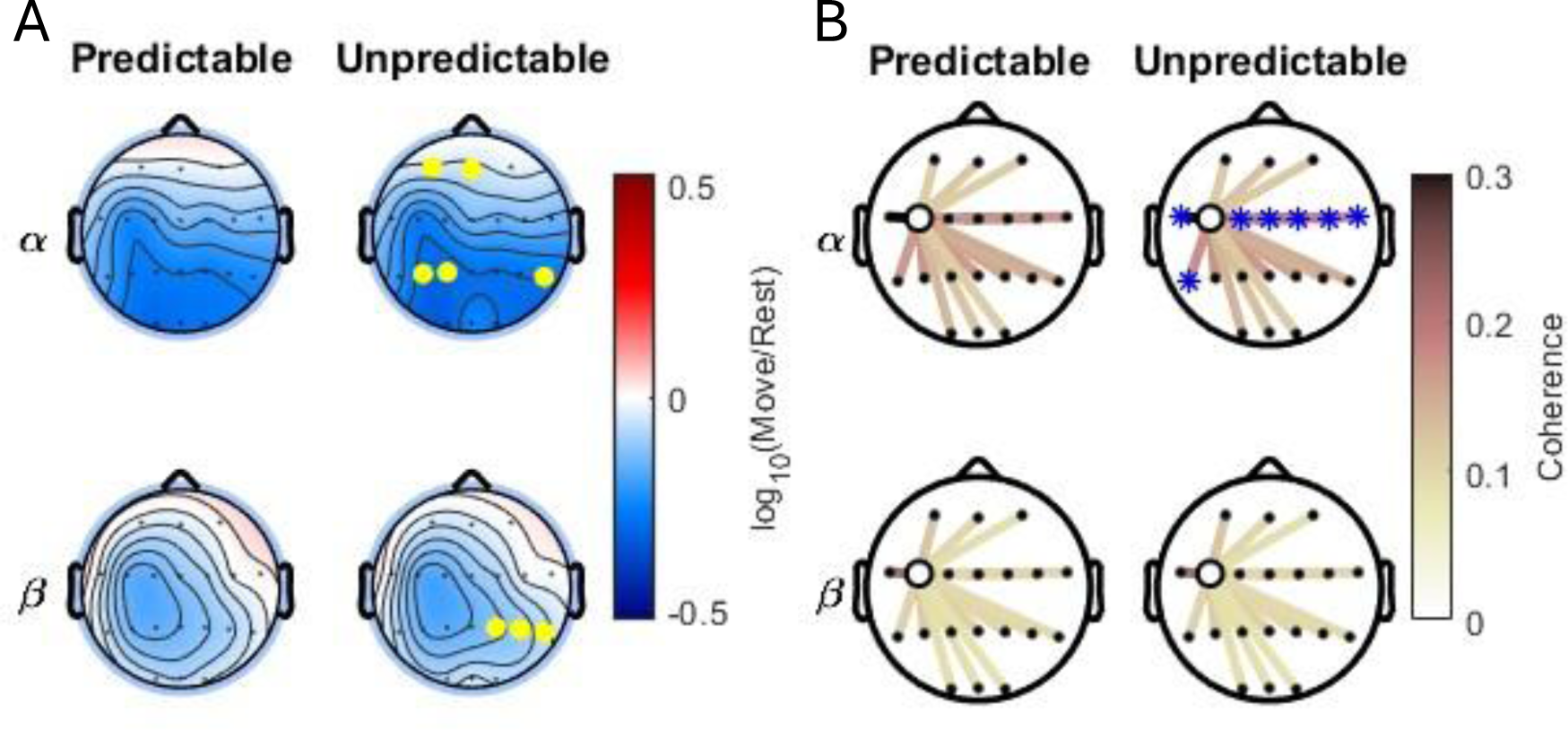
Scalp maps of the spectral features. (A) shows scalp maps of the ERDs averaged across the participants. Red and blue shades respectively indicate an increase or decrease in power when comparing rest to limb movements. Yellow circles indicate sensors with significantly stronger (p < 0.05) power reductions (i.e., ERDs) in the unpredictable condition compared to the predictable condition based on the paired-sample t-test. (B) shows scalp maps indicating the coherence values between other sensors and the bipolar sensor (FC3-C3). Darker graph lines indicate larger coherence values. Sensors marked with a blue asterisk indicate significantly larger (p < 0.05) coherence values in the unpredictable condition compared to the predictable condition based on the paired-sample t-test (N=20 participants).

Figure 4B shows graph plots of coherence between sensor FC3-C3 and the other sensors. Both conditions showed notably stronger alpha coherence values along the fronto-central sensors compared to other areas of the scalp. The alpha coherence values in these sensors were found to be significantly stronger in the unpredictable condition compared to the predictable condition: FC5-C5 (p = 0.003), FC1-C1 (p = 0.002), FCz-Cz (p < 0 0.001), FC2-C2 (p < 0.001), FC4-C4 (p = 0.005), FC6-C6 (p < 0.001), and CP5-P5 (p=0.012).

## Discussion

### Pursuing a target with unpredictable movements invokes higher levels of attention

In this study, we utilized a variant of the SART to assess how changes in attentional demand affected movement-related cortical activity as participants pursued a moving target. We found that participants were more accurate in correctly omitting key presses during the unpredictable condition. The unpredictable condition also yielded slower reaction times from the participants when they correctly responded to appropriate target shape. A higher accuracy with a slower reaction time is consistent with higher levels of attention from the participant, as interpreted in the SART protocol (Robertson et al., 1997). This matched our expectations that unpredictable movements from the target would require higher attention from participants as they pursued the target. While some aspects of this study are different from the conventional SART (e.g., utilizing shapes rather than numbers and setting the response omission target to appear 25% of the time), we found that it was sufficient to cause significant behavioral changes consistent with varying levels of attention.

### Frontal-parietal cortical activation and sensorimotor communication become stronger when motor tasks are performed with unpredictable perturbations

Based on the scalp maps of the ERDs, we found that the contralateral central areas had a notable power suppression in both the alpha and beta bands. For the alpha-band however, we also found substantial ERDs in the posterior areas of the scalp. This could be attributed to the nature of the task since tracking the target’s movement and shape likely induced a substantial engagement from the posterior areas associated with visual processing (Pfurtscheller et al., 1994). When we compared the ERDs between the two conditions however, we did not find a significant difference in the contralateral central areas. Instead, the alpha band ERDs were significantly stronger in the frontal and parietal areas while the beta band had significantly stronger ERDs in the ipsilateral parietal areas in the unpredictable condition. This reduction in alpha power during higher attention is consistent with previous neuroimaging studies that associate higher levels of engagement with weaker alpha band power (Jaquess et al., 2018; Pope et al., 1995).

Trials with unpredictable target movements also yielded significantly larger functional connectivity measures between the contralateral central sensor and the other sensors in the fronto-central areas across both hemispheres. This suggests that higher levels of attention yield stronger communication between other sensorimotor areas of the brain. The literature surrounding increased functional connectivity in sensorimotor areas during motor execution is limited but may provide context to the results in our study. Rilk and colleagues (2011) used a tracking task with a cursor that was controlled with grip force, and found that higher fronto-central coherence was correlated with higher tracking error. Manganotti and colleagues (1998) also found that bilateral fronto-central coherence was stronger when individuals performed more complex sequences of finger movement. Both factors (i.e., tracking error and task complexity) may have contributed to the results in this study, where the unpredictable condition with more complex movement trajectories resulted in higher tracking error.

Our results highlight that attention demanding tasks induce greater activation in the frontal and parietal areas while inducing greater synchrony in the sensorimotor areas. This could reflect top-down control signals from the prefrontal cortex and the posterior parietal cortex, which are thought to modulate activity in other cortical areas depending on attentional demands of the task (Ikkai & Curtis, 2011; Miller & Cohen, 2001; Seidler et al., 2012). Miller and Cohen (2001) proposed that the prefrontal cortex sends regulatory signals to the sensorimotor areas to activate preferential neural networks during motor practice; highlighting how the prefrontal cortex is engaged when individuals learn new motor skills and subsides when the motor skill is well-learned and performed with more automaticity. These top-down signals may be reflected in our results where higher attention was associated with stronger ERDs in the prefrontal areas, and the increased synchrony in the central areas could reflect the activation of preferential sensorimotor networks. Ikkai and Curtis (2011) proposed that the prefrontal cortex increases activity in the specific areas in the visual cortex that correspond to the individual’s attention to a receptive field. We anticipated an analogous result in our observed sensorimotor ERDs, but instead we only observed stronger connectivity in these regions. This contrast provides a new context of how attentional processes may modulate sensorimotor activity in the context of executing motor commands. At higher levels of attention during a motor task, the prefrontal cortex induces greater communication within the sensorimotor areas rather than strengthening activation. This increased communication could facilitate the formation of new neural connections that occur during motor learning. These results highlight a potential mechanism of how attention facilitates motor learning and stresses the importance of maintaining an individual’s attention during motor training and recovery.

It can be argued that Miller and Cohen’s (2001) hypothesis should be supported by an observation of larger coherence between the prefrontal areas and the sensorimotor areas during trials with unpredictable target movements. It is possible that the difference in coherence between the predictable and unpredictable conditions may have been attenuated due to a variety of factors. First, both conditions impose a high level of attention from the participants as they monitored changes in the target shape. Second, it is possible that the prefrontal cortex communicates with sensorimotor areas in a nonlinear fashion that would not be detected by coherence. Miller and Cohen (2001) also suggest that the prefrontal cortex sends regulatory signals that modulate sensorimotor activity, rather than simply passing similar signals between the two cortical areas. Lastly, although the prefrontal cortex also plays a large role in working memory, such processes would be less prevalent in this study since there were no cognitive demands related to memory recall or learning.

### Limitations

We intended to design the behavioral task such that only the level of attention was modulated between the two conditions. However, we also note that the tracking error between the cursor and the target was larger in the unpredictable condition. This indicates that the unpredictable condition was inherently more difficult, which may have contributed to the differences in the EEG features. Previous studies that used tracking tasks have found that frontal and parietal areas exhibit event-related potentials that are associated with tracking error (Krigolson et al., 2008; Krigolson & Holroyd, 2006). Alpha-band power has been found to be suppressed in frontal and parietal areas after participants make response errors in tasks like the Stroop Task (Carp & Compton, 2009, 2009) or the Simon Task (Driel et al., 2012). While we aimed for our results to reflect attentional processes, our results may be attributed to cognitive processes that perceive error in a motor task.

We also note that the results found in the study may reflect longitudinal learning or fatigue effects. All participants started with a block of predictable trials followed by the unpredictable trials. This study was designed this way because the dual task was demanding as it involved simultaneously performing two different tasks with different hands. Thus, we opted for participants to familiarize themselves with target movements that were easier and predictable before performing the task with unpredictable target movements. Nonetheless, the results of this study could be attributed to the order of the experimental conditions.

### Implications for motor learning and recovery

Our motivation behind this study was to elucidate how attention may interact with motor processes by directly observing movement-related neural activity. We can infer how these changes in neural activity can facilitate motor learning, but we stress that these inferences are limited to motor execution in the present study since the task was not a motor learning task. Nonetheless, we propose that emerging training and rehabilitative techniques could be enhanced based on the findings from this study.

Our study demonstrates that introducing random elements to a motor task can induce higher levels of attention, which may facilitate motor learning. While it is expected that participants would be more engaged in a task with unexpected changes in the environment, it is uncertain how this can be leveraged in programs for motor training and recovery. Adding unexpected perturbations to a training routine for the sake of maintaining attention is seldom done by trainers since it could confuse or discourage the trainee. However, our results support strategies that maintain attention from the trainee by adding variation to the practice. Training programs can be enhanced through contextual interference, which modifies practice schedules by alternating the practiced skills and/or the environment (Czyż et al., 2024; Shea & Morgan, 1979). Contextual interference has been known to result in better skill retention and skill transfer, even if the performance may be reduced due to the added variation (Shea & Morgan, 1979). We suspect these kinds of enhancements that add variation to a motor task like contextual interference could facilitate motor learning through the maintenance of attention. Our study provides a potential neural mechanism underlying the benefits of contextual interference and warrants further study to observe if contextual interference yields similar neural signatures related to attention and motor execution.

Our results also have implications for physical rehabilitation interventions that use biofeedback devices that monitor neural activity (Bhagat et al., 2020; Birbaumer et al., 2009; Donati et al., 2016; Irimia et al., 2017; Remsik et al., 2016; Venkatakrishnan et al., 2014). The motivation behind these systems is to encourage patients to be more engaged and attentive to rehabilitative exercises. These systems monitor the patient’s brain waves for their intent to move and provide positive feedback when intent is successfully detected. Such systems typically monitor movement-related features near sensorimotor areas of the scalp, such as alpha and beta-band ERDs (Buch et al., 2008; Fu et al., 2023; Irimia et al., 2017; Jeunet et al., 2019). However, our results imply that sensorimotor ERDs can have similar strengths at varying levels of attention. Our results highlight that incorporating prefrontal activity could provide more accurate assessments of the patient’s attention to the exercise and provides further support for rehabilitative neurofeedback systems that directly monitor the frontal area for patient engagement (Bartur et al., 2017; Rogers et al., 2021).

## Conclusion

In this study, we found that higher levels of attention in a target pursuit task was associated with stronger alpha band ERDs in the frontal and parietal areas, and stronger functional connectivity between the contralateral central areas with other sensors in the central areas of the scalp. While we anticipated that sensorimotor ERDs would be stronger with higher levels of attention, we found that they did not significantly change due to higher attentional demands. Our results concur with prevailing theories on attention and motor learning, where prefrontal processes related to attention may induce higher communication in sensorimotor areas. The neural signatures found in this study highlight a potential mechanism underlying how task variation can enhance motor learning. These neural signatures also provide a potential avenue to enhance rehabilitation techniques that utilize neurofeedback.

## Acknowledgements

This work was supported by the National Science Foundation under Grant # EEC-2127509 administrated by the American Society for Engineering Education (ASEE) to AYP.

## References

Ahn, S., Kim, K., & Jun, S. C. (2016). Steady-State Somatosensory Evoked Potential for Brain-omputer Interface—Present and Future. Frontiers in Human Neuroscience, 9. 10.3389/fnhum.2015.00716

Bartur, G., Joubran, K., Peleg-Shani, S., Vatine, J.-J., & Shahaf, G. (2017). An EEG Tool for Monitoring Patient Engagement during Stroke Rehabilitation: A Feasibility Study. BioMed Research International, 2017(1), 9071568. 10.1155/2017/9071568

Bartur, G., Joubran, K., Peleg-Shani, S., Vatine, J.-J., & Shahaf, G. (2020). A pilot study on the electrophysiological monitoring of patient’s engagement in post-stroke physical rehabilitation. Disability and Rehabilitation: Assistive Technology, 15(4), 471–479. 10.1080/17483107.2019.1680749

Bastos, A. M., & Schoffelen, J.-M. (2016). A Tutorial Review of Functional Connectivity Analysis Methods and Their Interpretational Pitfalls. Frontiers in Systems Neuroscience, 9. 10.3389/fnsys.2015.00175

Berka, C., Levendowski, D. J., Lumicao, M. N., Yau, A., Davis, G., Zivkovic, V. T., Olmstead, R. E., Tremoulet, P. D., & Craven, P. L. (2007). EEG Correlates of Task Engagement and Mental Workload in Vigilance, Learning, and Memory Tasks. 78(5).

Bhagat, N. A., Yozbatiran, N., Sullivan, J. L., Paranjape, R., Losey, C., Hernandez, Z., Keser, Z., Grossman, R., Francisco, G. E., O’Malley, M. K., & Contreras-Vidal, J. L. (2020). Neural activity modulations and motor recovery following brain-exoskeleton interface mediated stroke rehabilitation. NeuroImage: Clinical, 28, 102502. 10.1016/j.nicl.2020.102502

Bian, Y., Qi, H., Zhao, L., Ming, D., Guo, T., & Fu, X. (2018). Improvements in event-related desynchronization and classification performance of motor imagery using instructive dynamic guidance and complex tasks. Computers in Biology and Medicine, 96, 266–273. 10.1016/j.compbiomed.2018.03.018

Birbaumer, N., Ramos Murguialday, A., Weber, C., & Montoya, P. (2009). Chapter 8 Neurofeedback and Brain–Computer Interface: Clinical Applications. In International Review of Neurobiology (Vol. 86, pp. 107–117). Academic Press. 10.1016/S0074-7742(09)86008-X

Buch, E., Weber, C., Cohen, L. G., Braun, C., Dimyan, M. A., Ard, T., Mellinger, J., Caria, A., Soekadar, S., Fourkas, A., & Birbaumer, N. (2008). Think to Move: A Neuromagnetic Brain-Computer Interface (BCI) System for Chronic Stroke. Stroke, 39(3), 910–917. 10.1161/STROKEAHA.107.505313

Carp, J., & Compton, R. J. (2009). Alpha power is influenced by performance errors. Psychophysiology, 46(2), 336–343. 10.1111/j.1469-8986.2008.00773.x

Cassim, F., Szurhaj, W., Sediri, H., Devos, D., Bourriez, J.-L., Poirot, I., Derambure, P., Defebvre, L., & Guieu, J.-D. (2000). Brief and sustained movements: Differences in event-related (de)synchronization (ERD/ERS) patterns. Clinical Neurophysiology, 111(11), 2032–2039. 10.1016/S1388-2457(00)00455-7

Corbetta, M., & Shulman, G. L. (2002). Control of goal-directed and stimulus-driven attention in the brain. Nature Reviews Neuroscience, 3(3), 201–215. 10.1038/nrn755

Czyż, S. H., Wójcik, A. M., Solarská, P., & Kiper, P. (2024). High contextual interference improves retention in motor learning: Systematic review and meta-analysis. Scientific Reports, 14(1), 15974. 10.1038/s41598-024-65753-3

Delorme, A., & Makeig, S. (2004). EEGLAB: An open source toolbox for analysis of single-trial EEG dynamics including independent component analysis. Journal of Neuroscience Methods, 134(1), 9–21. 10.1016/j.jneumeth.2003.10.009

Donati, A. R. C., Shokur, S., Morya, E., Campos, D. S. F., Moioli, R. C., Gitti, C. M., Augusto, P. B., Tripodi, S., Pires, C. G., Pereira, G. A., Brasil, F. L., Gallo, S., Lin, A. A., Takigami, A. K., Aratanha, M. A., Joshi, S., Bleuler, H., Cheng, G., Rudolph, A., & Nicolelis, M. A. L. (2016). Long-Term Training with a Brain-Machine Interface-Based Gait Protocol Induces Partial Neurological Recovery in Paraplegic Patients. Scientific Reports, 6(1), 30383. 10.1038/srep30383

Driel, J. van, Ridderinkhof, K. R., & Cohen, M. X. (2012). Not All Errors Are Alike: Theta and Alpha EEG Dynamics Relate to Differences in Error-Processing Dynamics. Journal of Neuroscience, 32(47), 16795–16806. 10.1523/JNEUROSCI.0802-12.2012

Dujardin, K., Derambure, P., Defebvre, L., Bourriez, J. L., Jacquesson, J. M., & Guieu, J. D. (1993). Evaluation of event-related desynchronization (ERD) during a recognition task: Effect of attention. Electroencephalography and Clinical Neurophysiology, 86(5), 353.

Fu, J., Jiang, Z., Shu, X., Chen, S., & Jia, J. (2023). Correlation between the ERD in grasp/open tasks of BCIs and hand function of stroke patients: A cross-sectional study. Biomedical Engineering Online, 22(1), 36. 10.1186/s12938-023-01091-1

Giabbiconi, C. M., Dancer, C., Zopf, R., Gruber, T., & Müller, M. M. (2004). Selective spatial attention to left or right hand flutter sensation modulates the steady-state somatosensory evoked potential. Cognitive Brain Research, 20(1), 58–66. 10.1016/j.cogbrainres.2004.01.004

Hogan, N., Krebs, H. I., Rohrer, B., Palazzolo, J. J., Dipietro, L., Fasoli, S. E., Stein, J., Hughs, R., Frontera, W. R., Lynch, D., & Volpe, B. T. (2006). Motions or muscles? Some behavioral factors underlying robotic assistance of motor recovery. The Journal of Rehabilitation Research and Development, 43(5), 605. 10.1682/JRRD.2005.06.0103

Ikkai, A., & Curtis, C. E. (2011). Common neural mechanisms supporting spatial working memory, attention and motor intention. Neuropsychologia, 49(6), 1428–1434. 10.1016/j.neuropsychologia.2010.12.020

Irimia, D. C., Cho, W., Ortner, R., Allison, B. Z., Ignat, B. E., Edlinger, G., & Guger, C. (2017). Brain-Computer Interfaces With Multi-Sensory Feedback for Stroke Rehabilitation: A Case Study: BCI for Stroke Rehabilitation. Artificial Organs, 41(11), E178–E184. 10.1111/aor.13054

Jaquess, K. J., Lo, L.-C., Oh, H., Lu, C., Ginsberg, A., Tan, Y. Y., Lohse, K. R., Miller, M. W., Hatfield, B. D., & Gentili, R. J. (2018). Changes in Mental Workload and Motor Performance Throughout Multiple Practice Sessions Under Various Levels of Task Difficulty. Neuroscience, 393, 305–318. 10.1016/j.neuroscience.2018.09.019

Jeunet, C., Glize, B., McGonigal, A., Batail, J.-M., & Micoulaud-Franchi, J.-A. (2019). Using EEG-based brain computer interface and neurofeedback targeting sensorimotor rhythms to improve motor skills: Theoretical background, applications and prospects. Neurophysiologie Clinique, 49(2), 125–136. 10.1016/j.neucli.2018.10.068

Kahneman, D. (1973). Attention and effort. Prentice-Hall.

Krebs, H. I., Volpe, B., & Hogan, N. (2009). A working model of stroke recovery from rehabilitation robotics practitioners. Journal of NeuroEngineering and Rehabilitation, 6(1), 6. 10.1186/1743-0003-6-6

Krigolson, O. E., & Holroyd, C. B. (2006). Evidence for hierarchical error processing in the human brain. Neuroscience, 137(1), 13–17. 10.1016/j.neuroscience.2005.10.064

Krigolson, O. E., Holroyd, C. B., Van Gyn, G., & Heath, M. (2008). Electroencephalographic correlates of target and outcome errors. Experimental Brain Research, 190(4), 401–411. 10.1007/s00221-008-1482-x

Lelis-Torres, N., Ugrinowitsch, H., Apolinário-Souza, T., Benda, R. N., & Lage, G. M. (2017). Task engagement and mental workload involved in variation and repetition of a motor skill. Scientific Reports, 7(1), 14764. 10.1038/s41598-017-15343-3

Manganotti, P., Gerloff, C., Toro, C., Katsuta, H., Sadato, N., Zhuang, P., Leocani, L., & Hallett, M. (1998). Task-related coherence and task-related spectral power changes during sequential finger movements. Electroencephalography and Clinical Neurophysiology/Electromyography and Motor Control, 109(1), 50–62. 10.1016/S0924-980X(97)00074-X

Miller, E. K., & Buschman, T. J. (2013). Cortical circuits for the control of attention. Current Opinion in Neurobiology, 23(2), 216–222. 10.1016/j.conb.2012.11.011

Miller, E. K., & Cohen, J. D. (2001). An integrative theory of prefrontal cortex function. Annual Review of Neuroscience, 24, 167–202. 10.1146/annurev.neuro.24.1.167

Mizelle, J. C., Forrester, L., Hallett, M., & Wheaton, L. A. (2010a). Electroencephalographic reactivity to unimodal and bimodal visual and proprioceptive demands in sensorimotor integration. Experimental Brain Research, 203(4), 659–670. 10.1007/s00221-010-2273-8

Mizelle, J. C., Forrester, L., Hallett, M., & Wheaton, L. A. (2010b). Theta frequency band activity and attentional mechanisms in visual and proprioceptive demand. Experimental Brain Research, 204(2), 189–197. 10.1007/s00221-010-2297-0

Nakayashiki, K., Saeki, M., Takata, Y., Hayashi, Y., & Kondo, T. (2014). Modulation of event-related desynchronization during kinematic and kinetic hand movements. Journal of NeuroEngineering and Rehabilitation, 11(1), 90. 10.1186/1743-0003-11-90

Pfurtscheller, G. (2006). The cortical activation model (CAM). In Progress in Brain Research (Vol. 159, pp. 19–27). Elsevier. 10.1016/S0079-6123(06)59002-8

Pfurtscheller, G., & Lopes da Silva, F. H. (1999). Event-related EEG/MEG synchronization and desynchronization: Basic principles. Clinical Neurophysiology, 110(11), 1842–1857. 10.1016/S1388-2457(99)00141-8

Pfurtscheller, G., Neuper, C., & Mohl, W. (1994). Event-related desynchronization (ERD) during visual processing. International Journal of Psychophysiology, 16(2), 147–153. 10.1016/0167-8760(89)90041-X

Pfurtscheller, G., Stancák, A., & Neuper, Ch. (1996). Event-related synchronization (ERS) in the alpha band — an electrophysiological correlate of cortical idling: A review. International Journal of Psychophysiology, 24(1–2), 39–46. 10.1016/S0167-8760(96)00066-9

Pope, A. T., Bogart, E. H., & Bartolome, D. S. (1995). Biocybernetic system evaluates indices of operator engagement in automated task. Biological Psychology, 40(1), 187–195. 10.1016/0301-0511(95)05116-3

Remsik, A., Young, B., Vermilyea, R., Kiekhoefer, L., Abrams, J., Evander Elmore, S., Schultz, P., Nair, V., Edwards, D., Williams, J., & Prabhakaran, V. (2016). A review of the progression and future implications of brain-computer interface therapies for restoration of distal upper extremity motor function after stroke. Expert Review of Medical Devices, 13(5), 445–454. 10.1080/17434440.2016.1174572

Rilk, A. J., Soekadar, S. R., Sauseng, P., & Plewnia, C. (2011). Alpha coherence predicts accuracy during a visuomotor tracking task. Neuropsychologia, 49(13), 3704–3709. 10.1016/j.neuropsychologia.2011.09.026

Robertson, I. H., Manly, T., Andrade, J., Baddeley, B. T., & Yiend, J. (1997). ‘Oops!’: Performance correlates of everyday attentional failures in traumatic brain injured and normal subjects. Neuropsychologia, 35(6), 747–758. 10.1016/S0028-3932(97)00015-8

Rogers, J. M., Jensen, J., Valderrama, J. T., Johnstone, S. J., & Wilson, P. H. (2021). Single-channel EEG measurement of engagement in virtual rehabilitation: A validation study. Virtual Reality, 25(2), 357–366. 10.1007/s10055-020-00460-8

Seidler, R. D., Bo, J., & Anguera, J. A. (2012). Neurocognitive Contributions to Motor Skill Learning: The Role of Working Memory. Journal of Motor Behavior, 44(6), 445–453. 10.1080/00222895.2012.672348

Shea, J. B., & Morgan, R. L. (1979). Contextual interference effects on the acquisition, retention, and transfer of a motor skill. Journal of Experimental Psychology: Human Learning and Memory, 5(2), 179–187. 10.1037/0278-7393.5.2.179

Song, J.-H. (2019). The role of attention in motor control and learning. Current Opinion in Psychology, 29, 261–265. 10.1016/j.copsyc.2019.08.002

Veale, J. F. (2013). Edinburgh Handedness Inventory – Short Form: A revised version based on confirmatory factor analysis. Laterality, 19(2), 164–177. 10.1080/1357650X.2013.783045

Venkatakrishnan, A., Francisco, G. E., & L. Contreras-Vidal, J. (2014). Applications of Brain– Machine Interface Systems in Stroke Recovery and Rehabilitation. Current Physical Medicine and Rehabilitation Reports, 2(2), 93–105. 10.1007/s40141-014-0051-4

Wickens, C. D. (2020). Processing resources and attention. In D. L. Damos (Ed.), Multiple-task performance (1st ed., pp. 3–34). CRC Press. 10.1201/9781003069447-2

Yao, L., Meng, J., Zhang, D., Sheng, X., & Zhu, X. (2013). Selective Sensation Based Brain-Computer Interface via Mechanical Vibrotactile Stimulation. PLoS ONE, 8(6), e64784. 10.1371/journal.pone.0064784

Yuan, H., Perdoni, C., & He, B. (2010). Relationship between speed and EEG activity during imagined and executed hand movements. Journal of Neural Engineering, 7(2), 026001. 10.1088/1741-2560/7/2/026001

